# Chemical-microstructural-nanomechanical variations in the structural units of the cuttlebone of *Sepia officinalis*

**DOI:** 10.1101/156810

**Authors:** L North, David Labonte, ML Oyen, MP Coleman, HB Caliskan, RE Johnston

## Abstract

‘Cuttlebone’, the internalized shell found in all members of the cephalopod family Sepiidae, is a sophisticated buoyancy device combining high porosity with considerable strength. Using a complementary suite of characterization tools, we identified significant structural, chemical and mechanical variations across the different structural units of the cuttlebone: the dorsal shield consists of two stiff and hard layers with prismatic mineral organization which encapsulate a more ductile and compliant layer with a lamellar structure, enriched with organic matter. A similar organization is found in the lamellar matrix, which consists of individual chambers separated by septa, and supported by meandering plates (‘pillars’). Like the dorsal shield, septa contain two layers with lamellar and prismatic organization, respectively, which differ significantly in their mechanical properties: layers with prismatic organization are a factor of three stiffer, and up to a factor of ten harder than those with lamellar organization. The combination of stiff and hard, and compliant and ductile components may serve to reduce the risk of catastrophic failure, and reflect the role of organic matter for the growth process of the cuttlebone. Mechanically ‘weaker’ units may function as sacrificial structures, ensuring a step-wise failure of the individual chambers in cases of overloading, allowing the animals to retain near-neutral buoyancy even with partially damaged cuttlebones. Our findings have implications for the structure-property-function relationship of cuttlebone, and may help to identify novel bioinspired design strategies for light-weight yet high-strength foams.

## Introduction

Cuttlefish maintain near-neutral buoyancy at varying diving depths through the use of a specialized floatation device, frequently referred to as cuttlebone, which needs to combine high strength with minimum weight. Despite a porosity exceeding 90% (1), the cuttlebone of some species withstand pressures encountered at up to 500m in diving depth (2, 3). This exceptional combination of high compressive strength, porosity and permeability is extremely desirable for biomimetic and biomedical structural materials, including templates for tissue scaffolds (4, 5), hydroxyapatite scaffolds (6-8), and bone cements (9).

The remarkable performance of the cuttlebone is linked to its structural architecture: cuttlebone is composed of calcium carbonate (CaC0_3_) in its aragonite polymorph with a mixture of *β*-chitin and other protein complexes, and it comprises two main structural units: a dorsal shield and a lamellar matrix. The lamellar matrix is made up of continuous chambers, separated by parallel lamellae (septa), and supported by a complex arrangement of meandering plates (from here on referred to as ‘pillars’, see Fig. 1 A-D).

**Figure 1:**
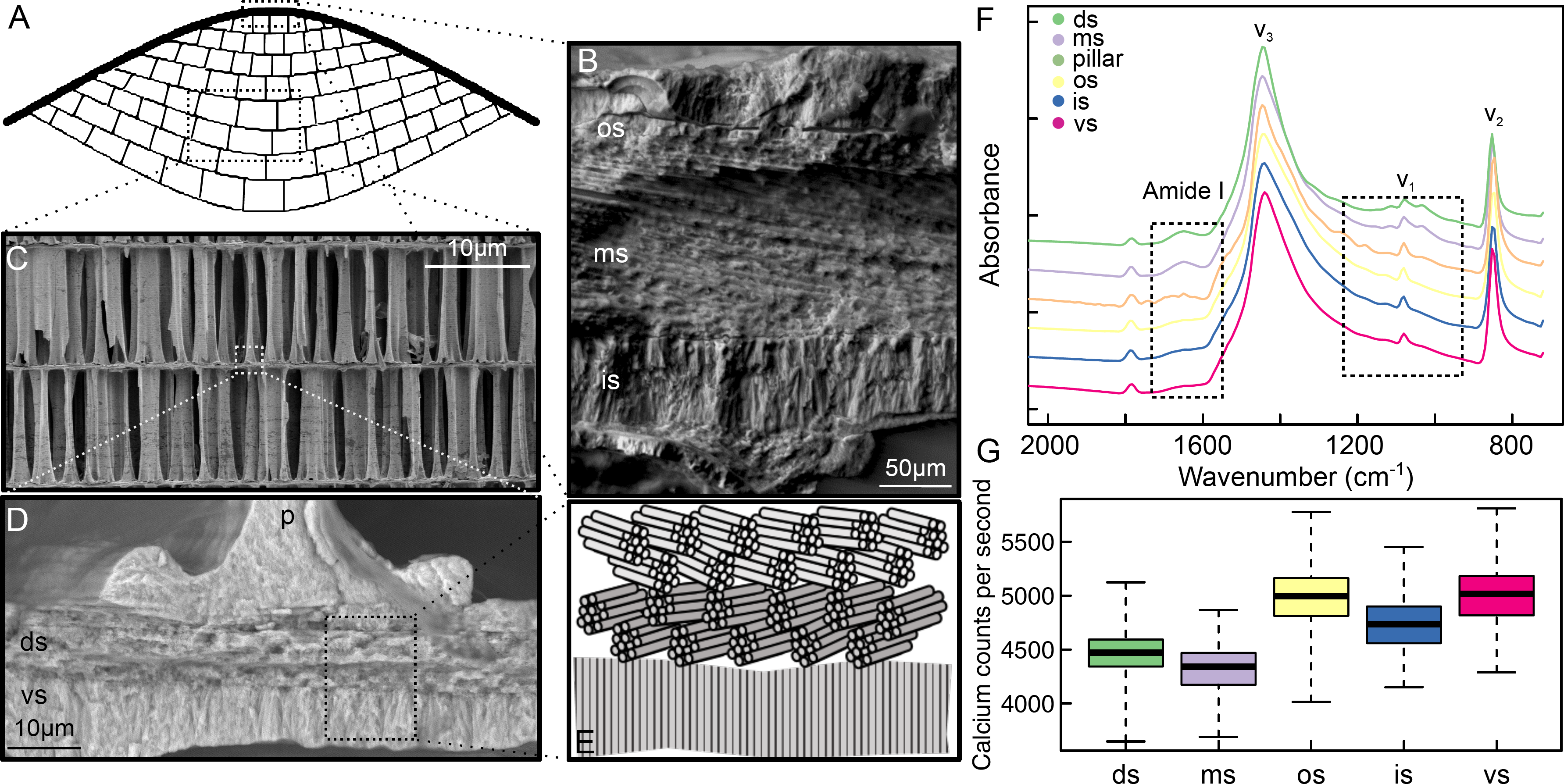
(A) Schematic cross-section of the cuttlebone, indicating the dorsal shield (thick black line) and lamellar matrix (cellular network, thin black lines), which consists of septa and pillars. Scanning electron micrographs of (B) the three distinct layers in the dorsal shield, (C) pillars supporting adjacent chambers, (D) the intersection of a septum and pillar illustrating the different microstructures within a septum. (E) Schematic of the variability in the orientation of fiber bundles in the septum. Chemical analyses of the distinct layered structures in cuttlebone in form of Attenuated total internal reflection Fourier Transform Infrared (ATR-FTIR) spectra are shown in (F). Each spectrum was normalized and then shifted vertically for clarity. Dashed boxes highlight the aragonite ‘fingerprint’ peak at 1080 cm^-1^ and the presence of organic macromolecules (chitin and possibly other proteins) at around 1650 cm^-1^, respectively. (G) Box-whisker plots illustrating the variation in calcium content between the three layers within the dorsal shield, and the two layers within septa, as measured with energy dispersive X-ray spectroscopy (EDS). Both FTIR and EDS reveal clear differences in the amount of organic matter and the Calcium content between the layers of both the dorsal shield and septa, which are similar among prismatic and lamellar crystal organizations, respectively. p - pillar, os - outer shield, ms - middle shield, is - inner shield, ds - dorsal septum and vs - ventral septum.

The macro-structure (1, 10-13), growth mechanisms (12, 14), bulk mechanical performance (1, 13), and chemical composition of cuttlebone (15), as well as the role of the organic constituents in the formation of the chambers and the organization of the mineralization have been studied in detail (1, 12, 14-16). However, the structure-property relationships across the identified structural units of this complex biomaterial have not yet been investigated, although they have been identified more than 180 years ago (17). In this work, we aim to close this gap, and provide a detailed chemical, structural and mechanical analysis of the different structural units, using scanning electron microscopy (SEM), energy dispersive X-ray spectroscopy (EDS), Fourier transform infrared spectroscopy (FTIR) and nanoindentation.

## Methods

Fresh specimens of *Sepia officinalis* (Linnaeus, 1758) were acquired commercially from around the UK coastline. The cuttlebone was dissected, sectioned into smaller samples (approximately 2 cm^3^) taken from the central region of the cuttlebone, and embedded in epoxy resin (EpoFix, Struers) using vacuum impregnation. Embedded specimens were diamond and colloidal silica polished to 0.04 μm finish, and a 7 nm carbon coating was applied using a Quorum Technologies Q150T turbo-pumped carbon coater.

EDS analysis in form of line scans at 10 kV was carried out on transverse sections to provide information on both the dorsal shield and lamellar matrix, using a Carl Zeiss Crossbeam 540 FEG SEM equipped with an SMax50 EDS detector, controlled via the Aztec software (Oxford Instruments). 10 line scans were performed with sampling separations within each line of 3.3and 0.14 μm for the dorsal shield and the septa, respectively; each line was measured three times and then averaged. The detailed number of spectra for individual structural units can be found in table S1 in the Supplemental Material.

Spectroscopic analysis of the samples was conducted using a PerkinElmer Spotlight400 Fourier transform infrared spectrometer (FTIR) fitted with an attenuated total internal reflection (ATR) imaging mode microscope (PerkinElmer, Waltham, Massachusetts, USA). The wavelength resolution was set to 12cm^-1^, and 8 scans per pixel were collected, with a pixel size of 1.56 μm.

Mechanical properties of the different structural units of the cuttlebone were measured with a TI 950 Triboindenter (Hysitron, Eden Prairie, USA), equipped with a diamond Berkovich indenter with a tip radius of 150 nm. Indentations were conducted in closed-loop displacement-control with a 5-10-5 s load-hold-unload trapezoidal load profile, and a peak displacement of 200 nm unless stated otherwise. Indentation hardness (H) and indentation modulus (E’) were measured from the force-displacement data using the Oliver & Pharr method (18). In order to quantify the changes in H and E’ across the shield, 18 horizontal lines of 25 vertical indents were placed across the shield of one specimen, starting on the septum adjacent to the ventral layer of the shield, and finishing on the surrounding epoxy. The vertical and horizontal spacings were 20 μm and at least 20 μm, respectively. The results were averaged either according to their vertical position (for plotting), or according to ‘region’ for further statistical analyses.

The mechanical properties of 14 randomly selected pillars were measured at various points along their length. The exact position varied across the pillars due to variable cross-sectional width, and we avoided regions that were thin or close to air gaps; the minimum horizontal spacing was 20 μm. A linear regression of *H* and *E’* against the indentation position normalized by pillar length revealed that neither differed significantly along the length of the pillars *(t* = 0.40, *p* = 0.69 and *t* = 0.61, *p* = 0.52 for *E’* and H, respectively). In order to capture the variation of material properties across pillars, a further 35 pillars were indented, with three indents per pillar, located approximately at the bottom, middle and top relative to the pillar length, and all results were pooled for further analyses.

Preliminary microscopy of the septum suggested that it contained at least two distinct regions (see Fig. 1 D-E), separated roughly at midline (see also 14, 16). In order to quantify the properties of these two regions, 60 indents were placed approximately centrally in each region, with a horizontal spacing of at least 20 μm.

In addition, we used the accelerated property-mapping (XPM) feature of the Triboindenter to visualize localized changes in material properties. XPM measurements were conducted in force-control, as recommended by the manufacturer. Mechanical properties of the shield were mapped with a 3 × 148 (3 μm horizontal × 3 μm vertical) grid, a peak load of 1 mN, and a 1-2-1 s trapezoidal load profile. The variation in mechanical properties at the transition between pillar and septum was visualized with a 12 × 25 (5 μm horizontal × 2 μm vertical) grid, with a peak load of 0.8 mN, a 1-5-1 s trapezoidal loading profile. Only measurements with a contact depth exceeding 60 nm were used for further analyses, and data excluded by this criterion were linearly interpolated from the surrounding data points. All XPM-data were used for plotting mechanical property maps and were not included in the quantitative statistical analysis.

## Results

The main structural components of the cuttlebone - the dorsal shield, the septa and the pillars - can be readily identified by light and electron microscopy (Fig. 1). The dorsal shield consists of two layers formed of prismatic tubercles that encapsulate a middle layer with lamellar aragonite structure (Fig. 1 B and 11). This difference is also apparent in backscattered electron imaging, where the sandwiched layer shows a reduced atomic density (see Supplementary Material Fig. S1 A). A similar structural arrangement was found for the septa, which are split at about midline (Fig. 1 D-E), confirming previous work which has shown that the dorsal part of the septa has an aragonite lamella-fibrillar microstructure with fiber bundles oriented in two different directions on different planes (14, 19). The prismatic structure of the ventral part, in turn, consists of aragonite fibers oriented parallel to the pillar height, which connect directly into the adjacent pillar (Fig. 1 D-E).

These structural differences were also evident in both FTIR spectra and EDS data. While the ‘fingerprint’ peaks of aragonite were present in all spectra, components with a lamellar ultrastructure showed a distinct broadening of the v_1_-mode peak (1080 cm^-1^), and additional low-intensity peaks between 1000 and 1300 cm^-1^, as well as at 1650^-1^, indicative of the presence of a significant amount of organic macromolecules, including *β*-chitin (17). Components with a prismatic morphology, in turn, had a higher Ca count per second (cps), as measured via EDS (4915 ± 16 compared to 4392 ± 11 cps, mean ± standard deviation, for detailed data on individual layers, see Fig. 1G and Supplementary Material Table S1). This difference translated into calcium weight percentages of 46.5% for the prismatic, and 45.15% for the lamellar components.

While these differences may sound minor, in combination they lead to a marked difference in mechanical properties. Nanoindentation revealed that units with prismatic structure have an indentation modulus which exceeds that of lamellar components by a factor between two and three, and an indentation hardness which is up to a factor of ten higher (see Figures 2 & 3 A-B), and Fig.4, and Supplemental Material Table S1 for detailed data). For both the dorsal shield and the septa, the changes in material properties between prismatic and lamellar building blocks appear to be abrupt and occurred on a microscopic length scale, as evident from XPM (see Fig 2 C & 3, respectively).

**Figure 2:**
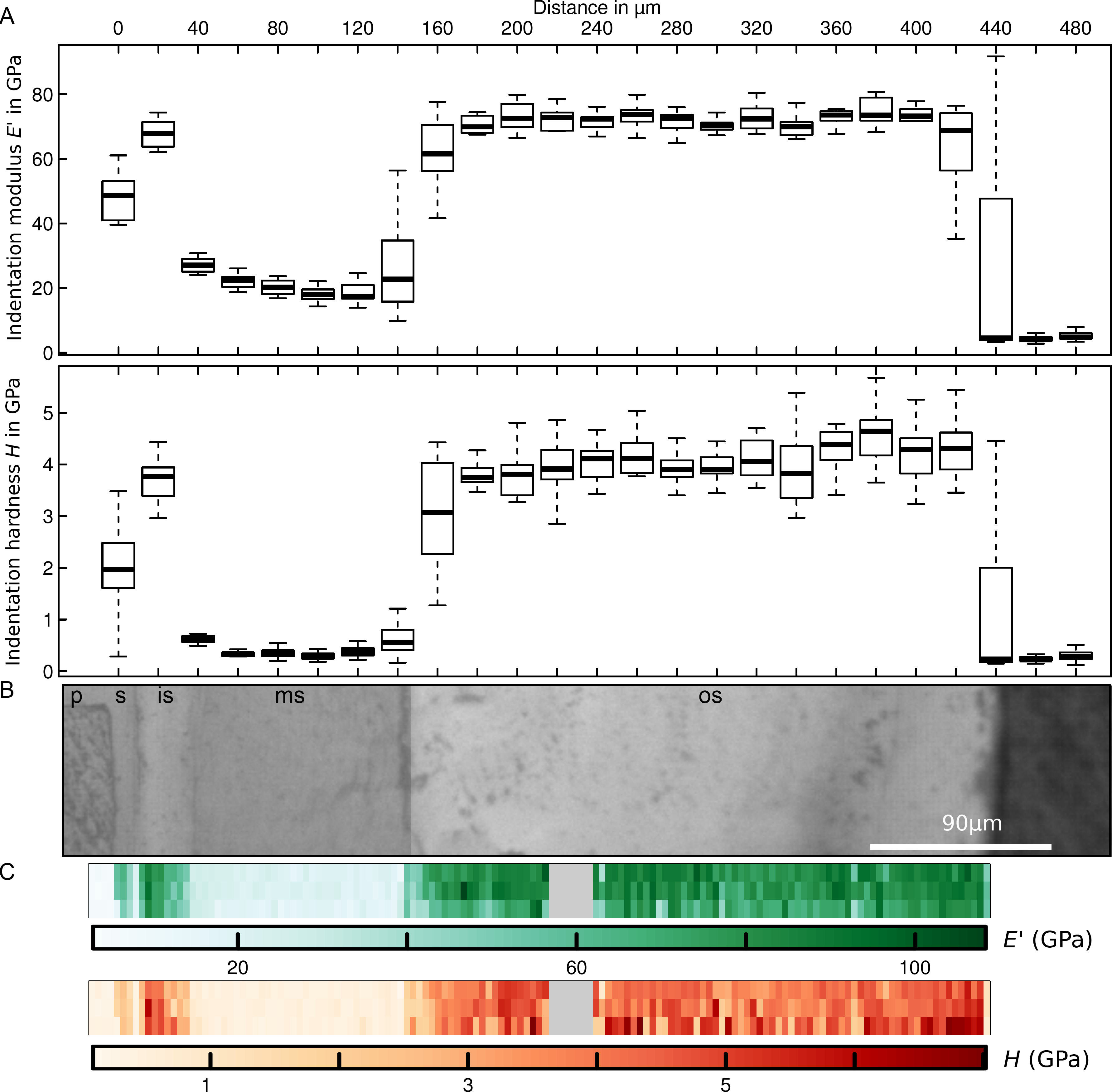
(A) Box-whisker plots illustrating the variation in indentation modulus and hardness across the different layers in the dorsal shield (n = 18 per box-whisker plot), which are shown in the light microscopy image in (B). The clear differences in indentation hardness and modulus and the apparent sharp transition between the different layers is visualized further in (C), which shows data collected with the accelerated property mapping feature of the 950 Triboindenter. The uniform block in (C) is a result of inaccuracies in the placement of the indents. p - pillar, os - outer shield, ms - middle shield, is - inner shield, s - septum.

**Figure 3:**
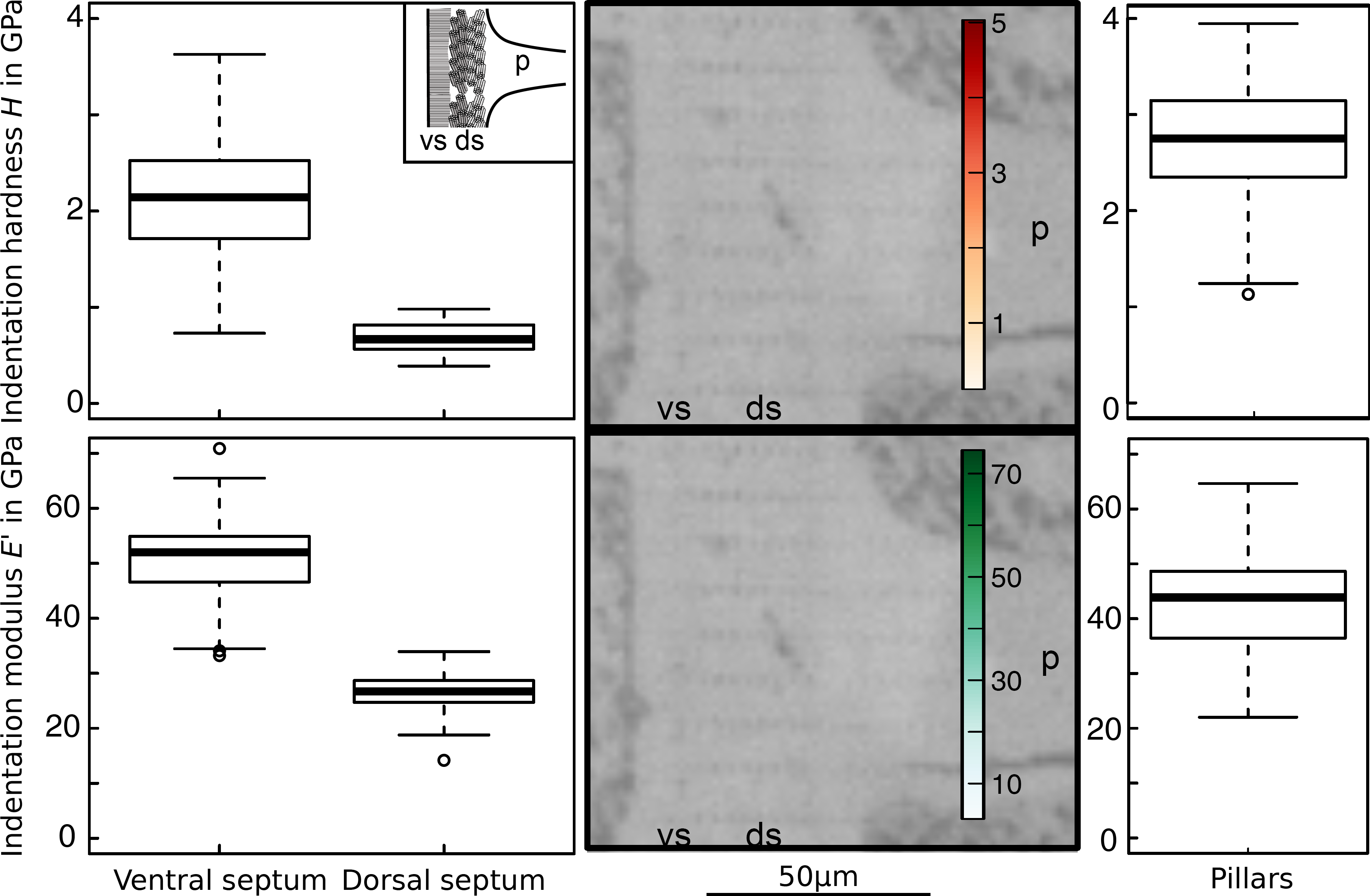
Box-whisker plots illustrating the variation in indentation modulus and hardness in the two distinct layers in the septa (left hand side, n = 60 per layer) and the pillars (right hand side, n = 275 for 49 different pillars). The clear differences in indentation hardness and modulus and the apparent sharp transition between the different layers is visualized further in the central region of the figure, which shows data collected with the accelerated property mapping feature of the 950 Triboindenter, overlaid on an SEM image of the corresponding region. P - pillar, ds - dorsal septum, vs - ventral septum.

## Discussion

Cuttlebone needs to be as light as possible to maximize buoyancy, but as strong as necessary to ensure sufficient safety factors. These conflicting demands are reflected in a complex bio-architecture, featuring at least six different structural units, which differ not only in their microstructure and chemical composition, but also in their mechanical properties (see Figs. 1-3): Stiff and hard layers with prismatic crystal organization alternate with more compliant and ductile zones, organized in a lamellar form and enriched with organic macromolecules. The inclusion of ‘weaker’ layers may at first seem surprising, as the maximum diving depth is directly related to the mechanical properties for a given geometrical arrangement (13, 20). However, several arguments may be brought forward to rationalize this design strategy.

Firstly, variations in indentation modulus between adjacent layers may represent a toughening mechanism. Cracks are effectively arrested at the interface between adjacent layers if the ratio of their elastic moduli is greater than about 5 (21), as for example observed in hexactinellid sponges, albeit at a much smaller scale (22). Both the middle layer of the dorsal shield and the dorsal septum might be particularly effective in arresting cracks, as they have an unusually small ratio between indentation hardness and modulus (about half the value of ≈ 0.05 typical for biological materials, see 23), which is a proxy for the tendency of a material to undergo elastic versus inelastic deformation (23). Hence, these layers may absorb energy by inelastic deformation instead of feeding it into the propagation of cracks. While these arguments are largely speculative at present, X-ray microCT images of the dorsal shield revealed several cases where cracks seemingly originating in the outer shield indeed failed to penetrate the central layer of the shield, and instead were arrested at the interface, providing some support for our hypothesis (see Supplementary Material Fig. S1 C).

Secondly, and related to the first point, the inclusion of weaker layers may represent a strategy to ensure controlled and localized structural failure in case of overloading, for example by directing gross failure to the pillars. The question remains, however, which failure mechanism would be most beneficial for the organism. Overloaded pillars collapse, resulting in a stepwise failure and densification of individual chambers (see Supplementary Material figure S1 B for a 4D X-ray microtomography visualization). Cuttlefish can survive multiple of such chamber failures (2, 3), and the variation in mechanical properties identified in this study might be key to avoid a damaging through-thickness crack in the dorsal shield, which would compromise near-neutral buoyancy.

Thirdly, the larger amount of organic molecules at the interface between adjacent chambers may arise as a necessary consequence of cuttlebone growth, as organic matter is known to control both nucleation and organization of biominerals (14, 16).

The way in which the cuttlebone fails when overloaded also has implications for the link between cuttlebone morphology and maximum diving depth established by earlier work (see e. g. 20). Previous studies have used beam or plate theory to study the mechanical performance of the *Sepia* cuttlebone with reasonable success (13, 20). While these studies assumed that failure occurs via elastic buckling of the pillars, there are two other competitive failure modes in foam-like solids, ‘plastic’ collapse or brittle crushing (24). Notably, the maximum sustainable stress, σ, predicted for the two latter failure modes is directly proportional to the aspect ratio characterizing the repetitive unit (24), which we define here as wall thickness, *t*, versus wall length, *l*. In contrast, 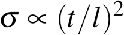 for elastic buckling (24), implying that elastic buckling will be the dominant failure mode for structures with small aspect ratios (see Supplementary Material). However, for animals which dive to deeper depths – and hence typically possess chambers with larger aspect ratios (20) – the mechanical performance limit might be set by plastic collapse or brittle crushing instead.

While the detailed functional implications of the identified variation in ultrastructure, chemistry and mechanical properties across the structural units of the cuttlebone require further experimental and numerical studies, this work will likely yield results not only relevant for our understanding of the structure-function relationship in the cuttlebone, but also for the design of bioinspired foams which combine low weight with high strength.

## Acknowledgements

The work was supported by the Advanced Imaging of Materials (AIM) facility (EPSRC Grant No. EP/M028267/1), the European Social Fund (ESF) through the European Union’s Convergence programme administered by the Welsh Government, the United States Army Corps of Engineers, Engineer Research and Development Centre (ERDC), the Denman Baynes Senior Research Fellowship (to DL), and an EPSRC Doctoral Training Award (EPSRC Grant No. EP/K502935/1 to LN). The authors would like to thank Peter Davies for microscopy technical assistance, and Dr Ed Pope for discussions on *Sepia officinalis* behavior.

## Appendix Failure modes of cuttlebone

We model the phragmacone as a regular foam with closed, cubic cells, characterised by an edge length *l*, and a wall thickness *t*, so that the relative density is 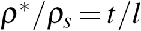 (see 24). Such a foam will collapse by elastic buckling at a critical stress σ*_el_* (24):

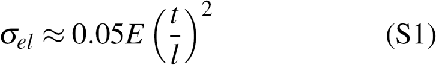

where we neglected the potential contribution arising from the compression of fluid inside the foam. Here, *E* is the Young’s modulus of the solid component of the foam. Alternatively, foams can fail by plastic collapse, at a critical pressure σ*_p_*, given by:

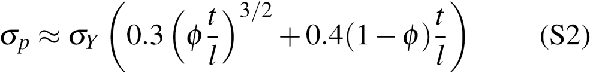

where σ*_Y_* is the yield strength, ϕ is the fraction of the solid contained in the cell edges (see below), and we again neglected the potential contribution arising from the compression of fluid inside the foam. Last, the foam can collapse by brittle crushing, at a critical stress defined by:

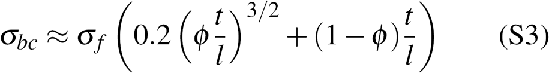

where σ_*f*_ is the modulus of rupture. For square prisms with constant wall thickness, 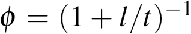, which is close to 0 for most phragmacones (20), so that the first term in eqs. 2-3 is much smaller than the linear term. As a consequence, the maximum stress is directly proportional to the aspect ratio for plastic collapse and brittle crushing, but proportional to the square of the aspect ratio for elastic buckling, which is the result referred to in the main manuscript.

---

**Table S1:** Summary of mechanical properties and calcium count per second for the different structural components of the cuttlebone investigated in this study. Data are given as mean ± standard deviation

|  | Indentation modulus E' | Indentation hardness H | H/E' | n | Calcium count |
| --- | --- | --- | --- | --- | --- |
| Outer shield | 71.8 ± 6.3 Gpa | 4.1 ± 0.6 Gpa | 0.056 ± 0.006 | 216 | 4991 ± 258 s<sup>-1</sup> |
| Middle shield | 20.0 ± 3.0 Gpa | 0.3 ± 0.1 Gpa | 0.017 ± 0.004 | 72 | 4320 ± 219 s<sup>-1</sup> |
| Inner shield | 68.0 ± 3.8 Gpa | 3.7 ± 0.4 Gpa | 0.054 ± 0.005 | 18 | 4748 ± 265 s<sup>-1</sup> |
| Pillars | 43.2 ± 9.3 Gpa | 2.7 ± 0.6 Gpa | 0.063 ± 0.012 | 275 | NA |
| Dorsal septum | 26.7 ± 3.4 Gpa | 0.7 ± 0.2 Gpa | 0.025 ± 0.004 | 60 | 4464 ± 196 s<sup>-1</sup> |
| Ventral septum | 51.0 ± 7.1 Gpa | 2.2 ± 0.6 Gpa | 0.041 ± 0.010 | 60 | 5007 ± 260 s<sup>-1</sup> |
| Epoxy | 4.7 ± 1.2 Gpa | 0.3 ± 0.1 Gpa | 0.056 ± 0.007 | 36 | NA |

**Figure S1:**
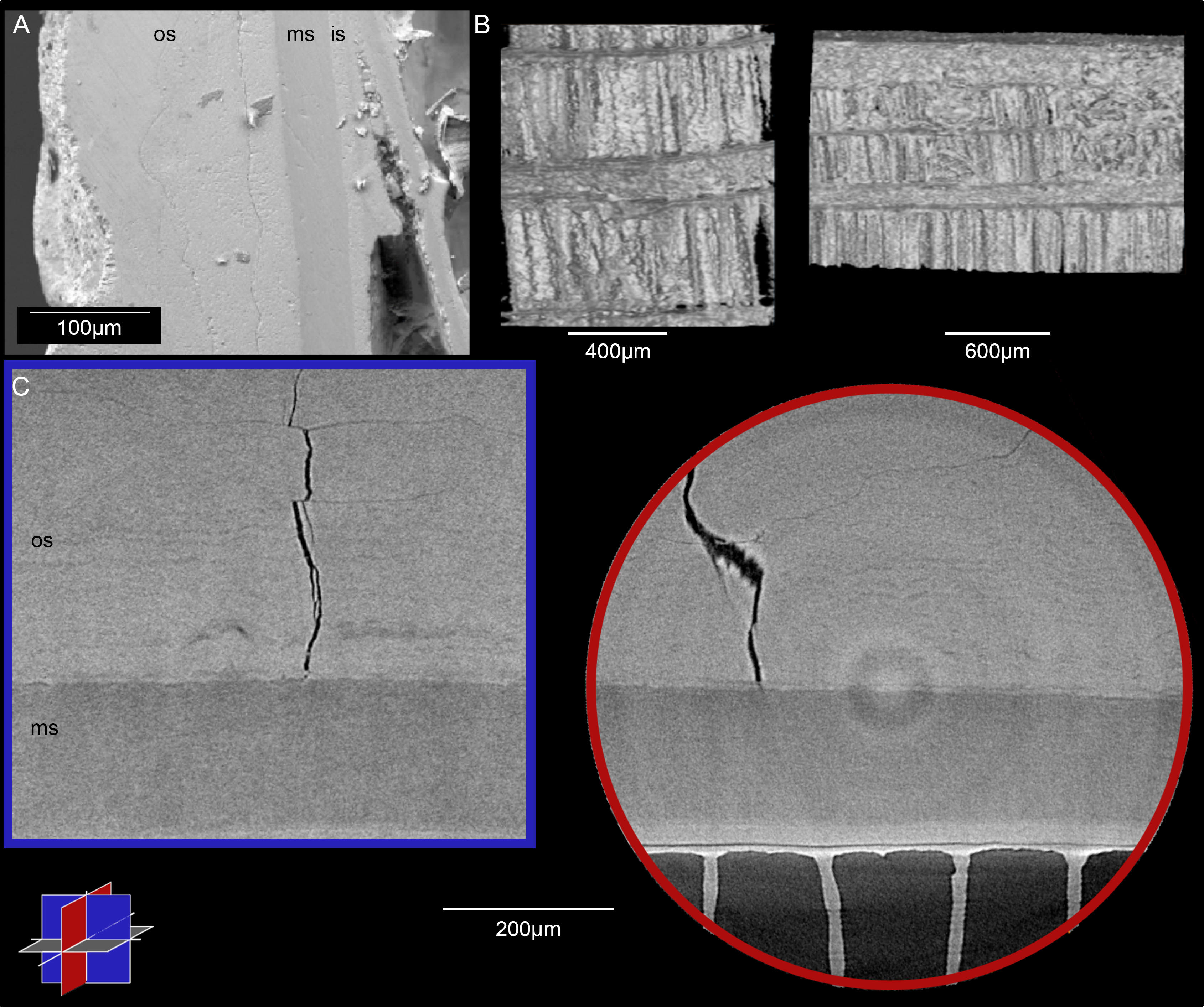
(A) Backscatter scanning electron micrograph of dorsal shield, identifying a layer of lower atomic mass (ms - middle shield) sandwiched between two layers of higher atomic mass (os - outer shield, is - inner shield). (B) X-ray microcomputed tomography of the failure in pillar structures within a layer and subsequent densification, visualized in three dimensions. (C) X-ray microcomputed tomography of the dorsal shield, visualised as 2D slices oriented perpendicular to each other, and to the plane of the shield. A crack can be seen extending through the uppermost layer, and failing to penetrate into the middle layer of the dorsal shield. The outer surface of the shield is oriented to the top of the slices.

